# Differential effects of cobalt ions *in vitro* on gill (Na^+^, K^+^)-ATPase kinetics in the blue crab *Callinectes danae* (Decapoda, Brachyura)

**DOI:** 10.1101/2022.11.18.516930

**Authors:** Francisco A. Leone, Leonardo M. Fabri, Maria I. C. Costa, Cintya M. Moraes, Daniela P. Garçon, John C. McNamara

## Abstract

To evaluate the crustacean gill (Na^+^, K^+^)-ATPase as a molecular marker for toxic contamination by heavy metals of estuarine and coastal environments, we provide a comprehensive analysis of the effects of Co^2+^ *in vitro* on modulation of the K^+^-phosphatase activity of a gill (Na^+^, K^+^)-ATPase from the blue crab *Callinectes danae*. Using *p*-nitrophenyl phosphate as a substrate, Co^2+^ can act as both stimulator and inhibitor of K^+^-phosphatase activity. Without Mg^2+^, Co^2+^ stimulates K^+^-phosphatase activity similarly but with a ≈4.5-fold greater affinity than with Mg^2+^. With Mg^2+^, K^+^-phosphatase activity is almost completely inhibited by Co^2+^. Substitution of Mg^2+^ by Co^2+^ slightly increases enzyme affinity for K^+^ and NH_4_^+^. Independently of Mg^2+^, ouabain inhibition is unaffected by Co^2+^. Mg^2+^ displaces bound Co^2+^ from the Mg^2+^-binding site in a concentration dependent mechanism. However, at saturating Mg^2+^ concentrations, Co^2+^ does not displace Mg^2+^ from its binding site even at elevated concentrations. Saturation by Co^2+^ of the Mg^2+^ binding site does not affect *p*NPP recognition by the enzyme. Given that the interactions between heavy metal ions and enzymes are particularly complex, their toxic effects at the molecular level are poorly understood. Our findings elucidate partly the mechanism of action of Co^2+^ on a crustacean gill (Na^+^, K^+^)-ATPase.

**Highlights:**

1. Without Mg^2+^, cobalt ions stimulate the gill (Na^+^, K^+^)-ATPase
2. Co^2+^ has a 4.5-fold greater affinity for the gill (Na^+^, K^+^)-ATPase than does Mg^2+^
3. Mg^2+^ displaces Co^2+^ from the Mg^2+^-binding site in a concentration dependent manner
4. Ouabain inhibition with Co^2+^ or Mg^2+^ is identical
5. Saturation by Co^2+^ of Mg^2+^-binding sites does not affect substrate recognition

**Graphical Abstract:** 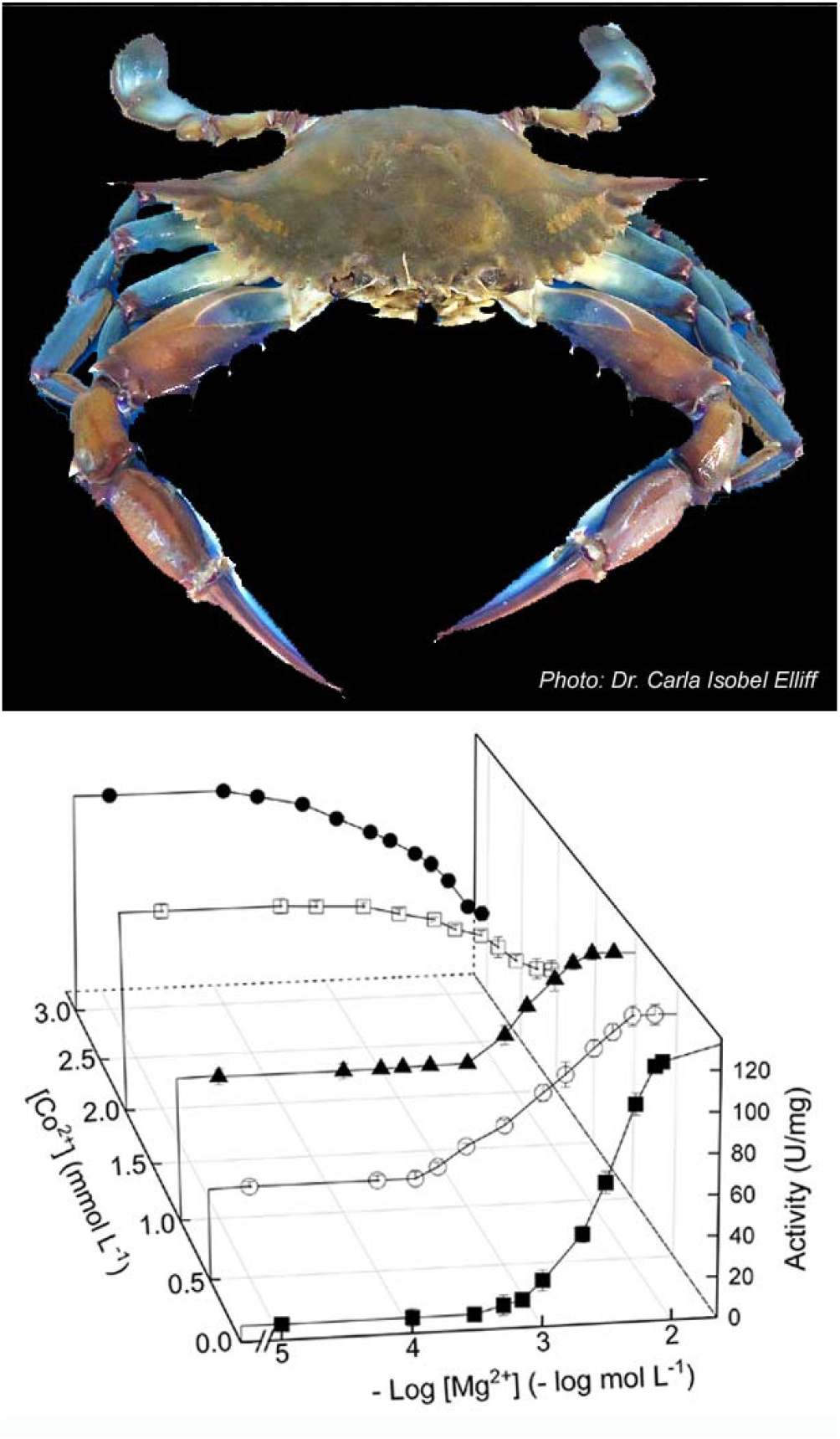

**Graphical abstract (synopsis):** Using a crab gill (Na^+^, K^+^)-ATPase, we demonstrate that Co^2+^ inhibits K^+^-phosphatase activity with Mg^2+^, which is stimulated without Mg^2+^. Mg^2+^ displaces Co^2+^ from the Mg^2+^-binding site but Co^2+^ cannot displace Mg^2+^. Ouabain inhibition is unaffected by Co^2+^, independently of Mg^2+^. The molecular mechanism of Co^2+^ toxicity is partly elucidated.

## 1. INTRODUCTION

Estuarine and coastal environments accumulate toxic contaminants owing to natural phenomena and/or anthropogenic activities [1]. Such pollutants include a wide variety of heavy metal ions, organic compounds and various micro/nano-particles [2–4]. Transition metals stand out specifically as the most abundant contaminants found in aquatic environments and contribute largely to toxicity [5,6]. Heavy metal ions are particularly harmful to organisms as they are not degradable and accumulate acutely or chronically in cells and tissues, altering biochemical and physiological homeostatic mechanisms, which can lead to demise at the organismal level [7].

The uptake and toxicity of heavy metals in aquatic organisms is determined by various ambient parameters like pH, temperature and salinity [8]. The biotic ligand model is the model most used to predict metal toxicity in aquatic environments [9]. A profusion of ecotoxicological studies has highlighted the importance of heavy metal toxicity in aquatic environments [e.g., 10,11,12,13]. Most studies of heavy metal toxicity in aquatic organisms concern their bioaccumulation in different tissues [1,14,15] and effects on physiological and biochemical processes like osmoregulatory capability and aerobic and oxidative stress metabolism [1,11,16]. The toxic effects of heavy metals at the molecular level are poorly known since the interactions between such metal ions and enzymes are complex [17,18]. The scant information available regarding the molecular mechanisms of heavy metal toxicity is limited mainly to fish and crustaceans [19–22].

While cobalt is a trace element indispensable for various physiological processes [21,23,24] it is toxic at high intracellular concentrations [25]. In humans, excessive exposure to cobalt results in complex health deficits involving heme oxidation, cytotoxicity, oxidative stress, apoptosis, altered membrane permeability, calcium channel blockage and DNA damage [26–28]. Active Ca^2+^ transport is inhibited in freshwater fish gills [22] while metabolic pathways are affected in freshwater algae [29].

Although cobalt concentrations in open oceanic waters are less than 7 ng L^−1^ [30], anthropogenic activities have led to progressive contamination of estuarine and coastal environments [see 25 for review]. To illustrate, mean cobalt levels measured in surface waters near an industrial plant in an Iberian Mediterranean estuarine system are ≈700 µg Co^2+^ L^−1^ and reach up to ≈2,800 µg Co^2+^ L^−1^ [31]. Cobalt titers in brachyuran crabs from marine/estuarine ecosystems range from 0.26 to 0.43 µg Co^2+^ g^−1^ soft tissue dry mass in *Callinectes sapidus* [32]; 0.91 to 1.2 µg Co^2+^ g^−1^ hepatopancreas dry mass in male and female *Portunus segnis* [33]; and 0.72 to 0.86 µg Co^2+^ g^−1^ dry mass in whole *Carcinus maenas* [34]. The lack of molecular information regarding harm to aquatic organisms, including the effects of Co^2+^, has impaired our overall comprehension of environmental damage and particularly of toxicity owing to bioaccumulation in marine organisms [16,23,25,35].

Estuarine and coastal organisms are powerful indicators of the health of the marine environment [36,37]. The portunid blue crab *Callinectes danae* is a crustacean species recommended as an environmental monitor of biological responses in contaminated estuarine and coastal areas [13]. The crab is a euryhaline osmoregulator, tolerant of exposure to wide range of salinities [38], and of great commercial value. It occurs from Florida (USA) to the southern Brazilian coast [39,40] and can be found in muddy estuaries and mangroves, on sandy and muddy shores, and in coastal waters up to 75 meters depth, including biotopes in which salinity varies from brackish to sea water [40,41].

In crustaceans, the gills provide a selective interface between the external environment and internal fluids, contributing to osmotic, ionic, excretory, and acid-base homeostasis. They are also an important route of entry of heavy metal ions [1,2,42]. Metal toxicity can thus vary depending on osmoregulatory strategy while alterations in salinity can affect the availability of metal ion species in the water column [36,43]. In brachyuran crabs, the three posterior gill pairs are specialized in ion transport [42], exhibiting a thickened epithelium [42,44] and increased expression and activity of ion transporters, including the (Na^+^, K^+^)-ATPase [1,45,46]. The presence of the (Na^+^, K^+^)-ATPase in the gill tissue and its role in physiological homeostasis renders this enzyme a suitable bioindicator to evaluate the kinetic effects of heavy metal ions like Co^2+^.

The (Na^+^, K^+^)-ATPase is a transmembrane enzyme that mediates the coupled transport of three Na^+^ from the cytosol into the extracellular fluid and of two K^+^ into the cytosol per ATP molecule hydrolyzed. The enzyme consists of two main subunits: a catalytic α-subunit, responsible for ion transport driven by ATP hydrolysis, and a highly glycosylated, non-catalytic β-subunit that modulates the transport properties of the enzyme [47,48]. The subunits are often associated with an FXYD peptide or γ-subunit that modulates (Na^+^, K^+^)-ATPase activity by altering the enzyme’s apparent affinity for Na^+^, K^+^ and ATP [49]. Briefly, the catalytic cycle involves alternation between the phosphorylated E1 and E2 conformations that show high affinity for Na^+^ and K^+^, respectively [47,48]. Magnesium is also essential although the ion is not transported during the catalytic cycle. Rather, Mg^2+^participates as the true substrate (a Mg•ATP complex) and plays a regulatory role, interfering with the conformational transitions of the enzyme [50,51]. At high concentrations, Mg^2+^ can inhibit Na^+^ and K^+^ binding by occupying a second specific inhibitory site outside the α-subunit membrane domains [50–52] or binding to the protein surface near the access channel of the ion binding sites [51,53,54]. Various divalent cations like Ca^2+^, Mn^2+^, Ba^2+^ and Sr^2+^ can substitute for Mg^2+^ by binding to the Mg^2+^ regulatory site, activating the enzyme [55–57]. The presence of both ATP and Na^+^ induces immediate enzyme phosphorylation such that Mg^2+^ binding and the phosphorylation reaction cannot be examined separately [51].

While the (Na^+^, K^+^)-ATPase exhibits high specificity for ATP it also catalyzes the ouabain-sensitive hydrolysis of other nucleoside triphosphates [58] and various non-nucleotide substrates such as *p*-nitrophenyl phosphate, acetyl phosphate, 2,4-dinitrophenyl phosphate, β-(2-furyl)-acryloyl phosphate, O-methyl fluorescein phosphate and 4-azido-2-nitrophenylphosphate [59–64]. The activity corresponding to the hydrolysis of non-nucleotide substrates is known as the K^+^-phosphatase activity and requires Mg^2+^ and K^+^ but is inhibited by Na^+^. Such activity represents a partial reaction of the (Na^+^, K^+^)-ATPase in which the E2 form is the main conformational state involved in *p*NPP hydrolysis [58]. The use of such non-nucleotide substrates has disclosed important kinetic characteristics of the (Na^+^, K^+^)-ATPase and, under most experimental conditions, these are better substrates than ATP itself [59,61–63,65]. Differently from ATP, *p*-nitrophenyl phosphate hydrolysis by the gill enzyme involves only a single substrate binding site [61,66]. Likewise, the (Na^+^, K^+^)-ATPase is also phosphorylated by *p*-nitrophenyl phosphate and other acyl phosphates [56,60,67,68]. K^+^ occlusion does not participate in phosphatase turnover [69] although, in the absence of Na^+^, K^+^ stimulates K^+^-phosphatase activity [70]. The use of *p*-nitrophenyl phosphate as a substrate thus facilitates the study of Mg^2+^ binding [51].

Given the paucity of information on the molecular effects of Co^2+^ on aquatic crustaceans from marine environments contaminated by heavy metals, in this study we provide a comprehensive analysis of the differential effects of Co^2+^ *in vitro* on the steady state kinetic properties of the K^+^-phosphatase activity of a gill (Na^+^, K^+^)-ATPase from the blue crab *Callinectes danae*.

## 2. MATERIAL AND METHODS

### Material

Millipore MilliQ (Merck KGaA, Darmstadt, Germany) ultrapure, apyrogenic deionized water was used to prepare all solutions. Chemicals of the highest purity commercially available were purchased from Sigma Chemical Co. (St. Louis, MO, USA) or Merck (Darmstadt, Germany). All salts were used as chlorides. The homogenization buffer consisted of 20 mmol L^−1^ imidazole (pH 6.8), 250 mmol L^−1^ sucrose and a proteinase inhibitor cocktail (1 mmol L^−1^ benzamidine, 5 µmol L^−1^ antipain, 5 µmol L^−1^ leupeptin,5 µmol L^−1^ phenyl-methane-sulfonyl-fluoride, and 1 µmol L^−1^ pepstatin A). Analytical estimation of the stock CoCl_2_ solution concentration (100 mmol L^−1^) was performed employing inductively coupled mass spectrometry (Perkin Elmer Avio 200 optical emission spectrometer, Boston MA, USA). NH_4_^+^ was removed from the crystalline ammonium sulfate suspensions of LDH and PK according to Fabri et al. [71]. When necessary, enzyme solutions were concentrated on YM-10 Amicon Ultra filters.

#### 2.1. Crab collection

Adult specimens of *Callinectes danae* of ≈9 cm carapace width were collected at low tide using seine nets or baited hand nets from Barra Seca beach (23° 25’ 01.5” S, 45° 03’ 01.0” W), Ubatuba, São Paulo State, Brazil (SISBIO/ICMBio/IBAMA authorization 02027.002342/98-04, permit #29594-18 to John C. McNamara). The crabs were held briefly in plastic boxes containing 30 L aerated seawater from the collection site during transport to the laboratory where they were immediately anesthetized by chilling in crushed ice. After bisecting and removal of the carapace, all three posterior gill pairs (120 gills/preparation, ≈5 g wet mass) were rapidly dissected out and frozen in liquid nitrogen in homogenization buffer in Falcon tubes.

#### 2.2. Preparation of the gill microsomal fraction

For each of the three (N= 3) gill homogenates prepared, the gills frozen in the homogenization buffer were thawed, diced and homogenized (20 mL homogenization buffer/g wet tissue) at 600 rpm in a Potter homogenizer in a crushed ice bath. The (Na^+^, K^+^)-ATPase-rich microsomal fraction was prepared by stepwise differential centrifugation (20,000 and 100,000 ×g) of the gill homogenate [38]. The resulting pellet was resuspended in 20 mmol L^−1^ imidazole buffer (pH 6.8) containing 250 mmol L^−1^ sucrose (15 mL buffer/g wet tissue). Aliquots (0.5 mL) were frozen in liquid nitrogen, stored at −20 °C and used within three-month’s storage provided that at least 95% (Na^+^, K^+^)-ATPase activity was present.

#### 2.3. Estimation of *p*-nitrophenyl phosphatase activity

*p*-Nitrophenyl phosphate (*p*NPP) was used as the enzyme substrate. The *p*-nitrophenyl phosphatase activity (*p*NPPase) of the microsomal fraction was estimated continuously at 25°C, following the release of the *p*-nitrophenolate ion (*p*NP^−^) at 410 nm (ε 410nm, pH 7.5= 13,160 M^−1^ cm^−1^) in a Shimadzu UV-1800 spectrophotometer (Vernon Hills IL, USA) equipped with thermostatted cells. The standard incubation medium contained 50 mmol L^−1^ HEPES buffer, pH 7.5, and appropriate *p*NPP, Mg^2+^, K^+^ or NH_4_^+^ concentrations (see Results and Figure legends for substrate and specific ionic concentrations) and 9 µg alamethicin, in final volume of 1 mL. Activity was estimated with or without 7 mmol L^−1^ ouabain, the difference corresponding to the K^+^-phosphatase activity of the gill (Na^+^, K^+^)-ATPase. The reaction was always initiated by the addition of the enzyme.

Because tissue homogenization usually results in the formation of small vesicles in the microsomal preparation that can occlude the catalytic site of the enzyme, *p*NPPase activity also was estimated after 10 min pre-incubation with 9 µg alamethicin, a membrane pore-forming antibiotic, to verify the presence of vesicles showing ATPase activity.

Controls without added enzyme were used to estimate the spontaneous hydrolysis of the substrate under the assay conditions. The kinetic measurements were carried out in duplicate, and substrate hydrolysis was accompanied over the shortest possible period to guarantee initial velocity measurements (<5% of substrate hydrolyzed during the reaction period). One unit (U) of enzyme activity was defined as the amount of enzyme that hydrolyses 1.0 nmol of *p*NPP per minute at 25 °C.

#### 2.4. Protein

Protein concentration was measured in duplicate aliquots of the microsomal preparations using the Coomassie Blue G dye-binding assay employing bovine serum albumin as a standard. Assays were read at 595 nm using a Shimadzu UV-1800 spectrophotometer [71].

#### 2.5. Estimation K^+^-phosphase activity with cobalt ions

The effect of cobalt ions on *p*NPPase activity was estimated as above using cobalt concentrations between 10^−5^ and 2×10^−2^ mmol L^−1^. Our study on the effect of Co^2+^ on the interaction of the different ligands with the (Na^+^,K^+^)-ATPase was performed using 3 mmol L^−1^ Co^2+^ (50.8 μg L^−1^ Co^2+^) which inhibits K^+^-phosphatase activity by ≈50%.

#### 2.6. Measurement of ATP hydrolysis

To provide a direct comparison with previous microsomal preparations, initial rates of ATP hydrolysis also were estimated continuously at 25 °C, using a pyruvate kinase/lactic dehydrogenase coupling system in which ATP hydrolysis is coupled to NADH oxidation [72]. The specific activity of (Na^+^, K^+^)-ATPase for ATP of 296.5 ± 10.2 nmol Pi min^−1^ mg^−1^ protein corresponding to 125.5 ± 2.3 nmol *p*NP^−^ min^−1^ mg^−1^ protein of K^+^-phosphatase activity is comparable to our previous findings [73].

#### 2.7. Estimation of kinetic parameters

The kinetic parameters V_M_ (maximum velocity), K_0.5_ (apparent dissociation constant) and the n_H_ value (Hill coefficient) for *p*NPP hydrolysis were calculated using SigrafW software ([74]; freely available from http://portal.ffclrp.usp.br/sites/fdaleone/downloads). The apparent dissociation constant of the enzyme-inhibitor complex (K_I_) was estimated using the Dixon plot in which the reaction rate corresponding to the K^+^-phosphatase activity was corrected for residual activity at high inhibitor concentrations [75]. Data points (mean ± SD) in the figures representing each substrate/ligand concentration are mean values of duplicate aliquots from the same preparation and were used to fit the saturation curves that were repeated using three different microsomal preparations (N= 3). The kinetic parameters (V_M_, K_M_, K_I_ or K_0.5_) shown in the tables are calculated values and represent the mean (± SD) derived from values estimated for each of three (N= 3) microsomal preparations.

## 3. RESULTS

Estimation of *p*NPPase activity with alamethicin revealed that the gill microsomal preparation from fresh caught *C. danae* includes ≈25% sealed, *p*NPPase-containing vesicles (164.5 ± 3.8 and 122.5 ± 1.8 nmol *p*NP^−^ min^−1^ mg^−1^ protein with or without alamethicin, respectively). Thus, *p*NPPase activity was always estimated using 9 µg alamethicin. Seven mmol L^−1^ ouabain decreased *p*NPPase activity from 164.5 ± 3.8 to 39.2 ± 1.9 nmol *p*NP^−^ min^−1^ mg^−1^ protein, indicating that ≈75% of the total phosphohydrolyzing activity (125.5 ± 2.3 nmol *p*NP^−^ min^−1^ mg^−1^ protein) corresponds to the K^+^-phosphatase activity of the gill (Na^+^, K^+^)-ATPase. Measurements using ATP as a substrate showed a (Na^+^, K^+^)-ATPase activity of 296.5 ± 10.2 nmol Pi min^−1^ mg^−1^ protein.

### 3.1. Effect of Co^2+^ on K^+^-phosphatase activity

Under optimal assay conditions (see legends to Fig. 1A and 1B), increasing Co^2+^ concentrations from 10^−5^ to 2×10^−2^ mol L^−1^ in the incubation medium inhibited K^+^-phosphatase activity by 90% (Fig. 1A and Table 1) with K_I_= 2.77 ± 0.33 mmol L^−1^ (inset a to Fig. 1). K^+^-phosphatase activity decreased following a single titration curve to 13.7 ± 3.6 nmol pNP^−^ min^−1^ mg^−1^ protein. The ouabain-insensitive *p*NPPase activity of ≈38 nmol pNP^−^ min^−1^ mg^−1^ protein was unaffected by increasing Co^2+^ concentrations (inset b to Fig. 1). On fixing Co^2+^ at 3 mmol L^−1^, increasing Mg^2+^ concentrations (10^−5^ to 2×10^−2^ mol L^−1^) inhibited K^+^-phosphatase activity by ≈60% (Fig. 1B and Table 1). A K_I_= 4.81 ± 0.71 mmol L^−1^ was calculated for inhibition of K^+^-phosphatase activity by Mg^2+^ in the presence of Co^2+^ (inset to Fig. 1B).

**Table 1.**
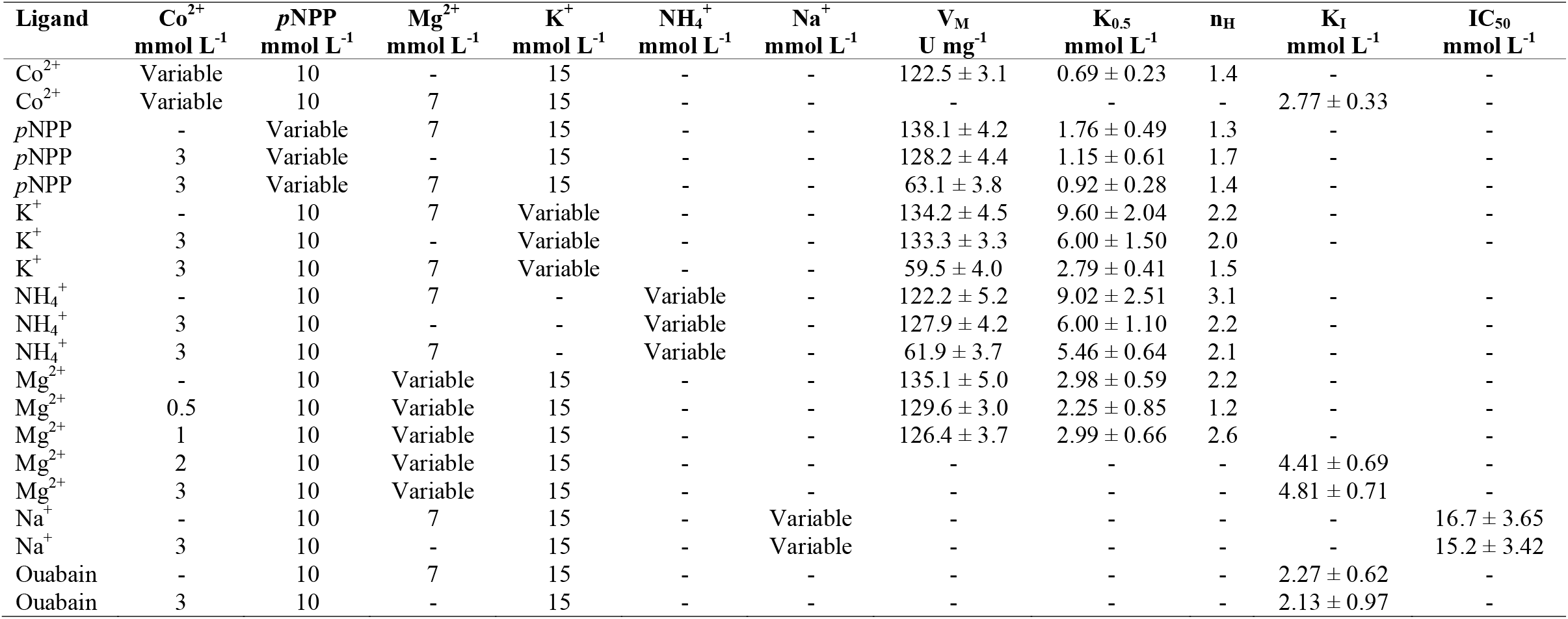
Kinetic parameters calculated for the effect of *p*NPP, Mg^2+^, K^+^, NH_4_^+^, Na^+^, Co^2+^ and ouabain on K^+^-phosphatase activity of (Na^+^, K^+^)-ATPase in a gill microsomal preparation from *Callinectes danae*.

**Figure 1.**
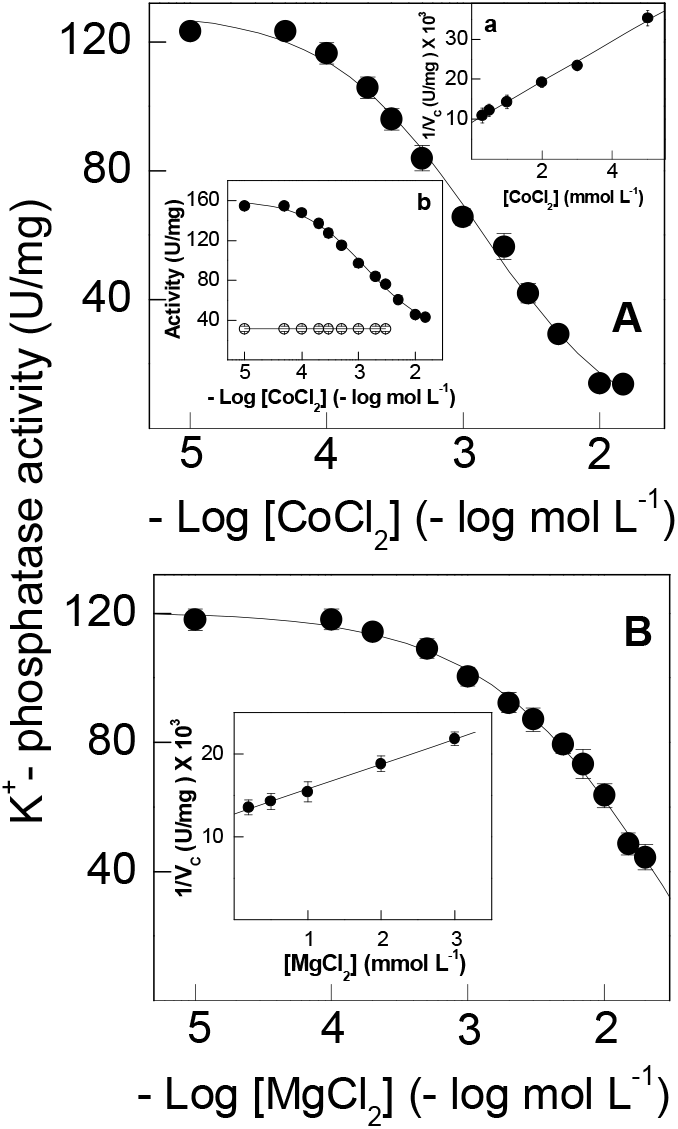
Effect of cobalt or magnesium ions in the presence of each other on the K^+^-phosphatase activity of *C. danae* gill (Na^+^, K^+^)-ATPase. Activity was assayed continuously at 25 °C in 50 mmol L^−1^ HEPES buffer, pH 7.5, containing 10 mmol L^−1^ *p*NPP, 15 mmol L^−1^ KCl, 9 µg alamethicin and the metal ions in a final volume of 1 mL. The mean activity of duplicate aliquots of the same microsomal preparation (≈15 µg protein) was used to fit the saturation curve which was repeated using three different microsomal preparations (± SD, N= 3). Where lacking, error bars are smaller than the symbols used. **A-** with 7 mmol L^−1^ MgCl_2_. Inset a**-** Dixon plot for estimation of K_I_ in which v_c_ is the K^+^-phosphatase activity corrected for residual pNPPase activity found at high Co^2+^ concentration. Inset b**-** total pNPPase activity (•) and ouabain-insensitive pNPPase activity (○). **B-** with 3 mmol L^−1^ CoCl_2_. Inset**-** Dixon plot for estimation of K_I_ in which v_c_ is the K^+^-phosphatase activity corrected for residual K^+^-phosphatase activity found at high Mg^2+^ concentrations.

Cobalt ions can substitute for Mg^2+^, stimulating gill K^+^-phosphatase activity (Fig. 2A and Table 1). Under optimal assay conditions (see legends to Fig. 2A and 2B) without Mg^2+^, Co^2+^ stimulated K^+^-phosphatase activity to a maximum rate of V_M_= 122.5 ± 3.1 nmol *p*NP^−^ min^−1^ mg^−1^ protein with K_0.5_= 0.69 ± 0.23 mmol L^−1^, showing a single saturation curve obeying cooperative kinetics (n_H_= 1.4). Ouabain-insensitive *p*NPPase activity was stimulated to 46.4 ± 3.7 nmol *p*NP^−^ min^−1^ mg^−1^ protein over the same Co^2+^ concentration range (inset to Fig. 2A). K^+^-phosphatase activity was inhibited by Co^2+^ concentrations above 10^−2^ mol L^−1^. In the absence of Co^2+^, Mg^2+^ (10^−4^ to 2×10^−2^ mol L^−1^) stimulated K^+^-phosphatase activity to a maximum rate of V_M_= 135.1 ± 5.0 nmol *p*NP^−^ min^−1^ mg^−1^ protein with K_0.5_= 2.98 ± 0.59 mmol L^−1^ and cooperative kinetics (n_H_= 2.2), following a single saturation curve (Fig. 2B and Table 1). Stimulation by Mg^2+^ of ouabain-insensitive *p*NPPase activity was negligible over the concentration range used (inset to Fig. 2B). The ≈4.5-fold lower K_0.5_ suggests that Co^2+^ binds more tightly to the Mg^2+^-binding sites than does Mg^2+^ itself.

**Figure 2.**
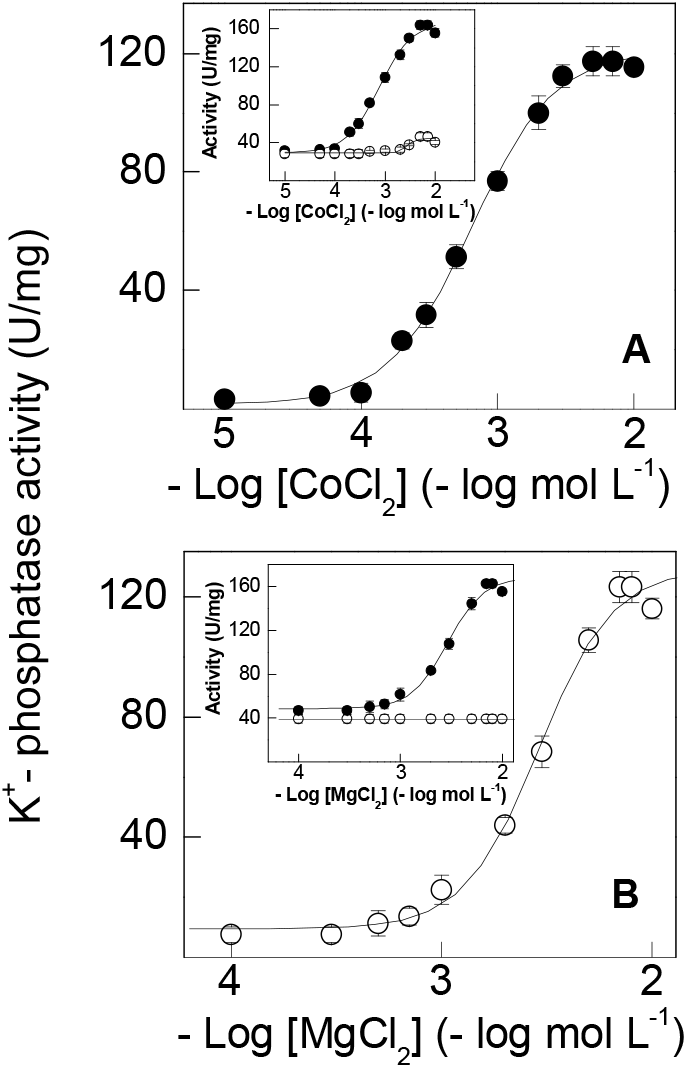
Estimulation by cobalt or magesium ions of K^+^-phosphatase activity of *C. danae* gill (Na^+^, K^+^)-ATPase. Activity was assayed continuously at 25 °C in 50 mmol L^−1^ HEPES buffer, pH 7.5, containing 10 mmol L^−1^ *p*NPP, 15 mmol L^−1^ KCl, 9 µg alamethicin in a final volume of 1 mL. The mean activity of duplicate aliquots of the same microsomal preparation (≈15 µg protein) was used to fit the saturation curve which was repeated using three different microsomal preparations (± SD, N= 3). Where lacking, error bars are smaller than the symbols used. **A-** with cobalt ions. **B-** with magnesium ions. Inset to figures-total *p*NPPase activity (•); ouabain-insensitive *p*NPPase activity (◯).

### 3.2. Effect of Co^2+^ on pNPP hydrolysis

Under optimal assay conditions (see legends to Fig. 2A and 2B), increasing *p*NPP concentrations stimulated K^+^-phosphatase activity to a maximum rate of V_M_= 138.1 ± 4.2 nmol *p*NP^−1^ min^−1^ mg^−1^ protein with K_0.5_= 1.76 ± 0.49 mmol L^−1^, following cooperative kinetics (n_H_= 1.3) and showing a single saturation curve (Fig. 3A and Table 1). Substitution of Mg^2+^ with 3 mmol L^−1^ Co^2+^ also gave a single saturation curve, overlapping that for Mg^2+^, showing a maximum rate of V_M_= 128.2 ± 4.4 nmol *p*NP^−^ min^−1^ mg^−1^ protein with K_0.5_= 1.15 ± 0.61 mmol L^−1^ (Fig. 3A). With Mg^2+^, the ouabain-insensitive *p*NPPase activity of ≈30 nmol *p*NP^−^ min^−1^ mg^−1^ protein was not affected by increasing *p*NPP concentrations (not shown). However, with 3 mmol L^−1^ Co^2+^ this activity was stimulated by ≈35% over the same *p*NPP concentration range (inset to Fig. 3A). With both Mg^2+^ and Co^2+^, under the same saturating ionic and substrate concentrations, K^+^-phosphatase activity decreased to a maximum rate of V_M_= 63.1 ± 3.8 nmol *p*NP^−^ min^−1^ mg^−1^ protein with K_0.5_= 0.92 ± 0.28 mmol L^−1^ also obeying cooperative kinetics (Fig. 3B and Table 1). This ≈50% inhibition of K^+^-phosphatase activity was accompanied by a 2-fold decrease in K_0.5_ compared to Mg^2+^ (Table 1). The ouabain-insensitive *p*NPPase activity of ≈20 nmol *p*NP^−^ min^−1^ mg^−1^ protein was stimulated ≈60% over the same *p*NPP concentration range (inset to Fig. 3B).

**Figure 3.**
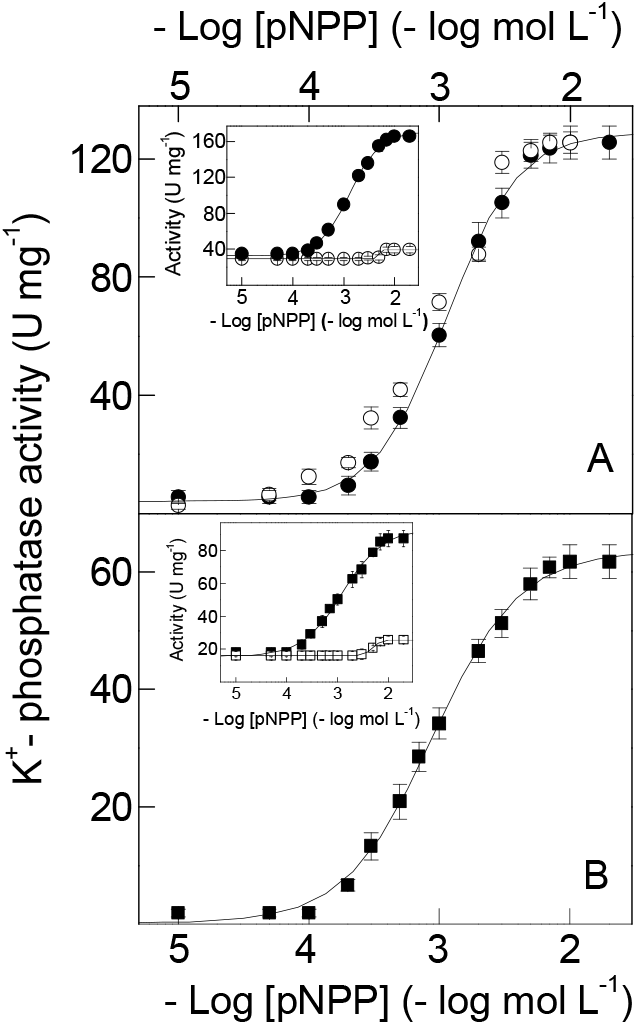
Effect of cobalt or magnesium ions on the modulation by *p*NPP of the K^+^-phosphatase activity of *C. danae* gill (Na^+^, K^+^)-ATPase. Activity was assayed continuously at 25 °C in 50 mmol L^−1^ HEPES buffer, pH 7.5, containing 15 mmol L^−^1 KCl, 9 µg alamethicin and the metal ion (7 mmol L^−1^ Mg^2+^ or 3 mmol L^−1^ Co^2+^) in a final volume of 1 mL. The mean activity of duplicate aliquots of the same microsomal preparation (≈15 µg protein) was used to fit the saturation curve which was repeated using three different microsomal preparations (± SD, N= 3). Where lacking, error bars are smaller than the symbols used. **A-** with magnesium (○) or cobalt ions (•**). B-** with both magnesium and cobalt ions. **Inset to figures-** total *p*NPPase activity (•,◼); ouabain-insensitive *p*NPPase activity (○,□) for Co^2+^only.

### 3.3. Effect of Co^2+^ on Mg^2+^ stimulation

Magnesium can displace Co^2+^ bound to the gill (Na^+^, K^+^)-ATPase below 2 mmol L^−1^ Co^2+^ (Fig. 4). Increased Mg^2+^ (10^−5^ to 5×10^−2^ mol L^−1^) displaces bound Co^2+^ (0.5 and 1 mmol L^−1^), stimulating K^+^-phosphatase activity to a maximum rate of ≈130 nmol *p*NP^−^ min^−1^ mg^−1^ protein with similar K_0.5_ (Fig. 4 and Table 1). However, at 2 mmol L^−1^ Co^2+^ and above no displacement was seen and increasing Mg^2+^ concentrations inhibited K^+^-phosphatase activity with K_I_= 4.41 ± 0.69 and 4.81 ± 0.71 mmol L^−1^ for 2 and 3 mmol L^−1^ Co^2+^, respectively (Table 1).

**Figure 4.**
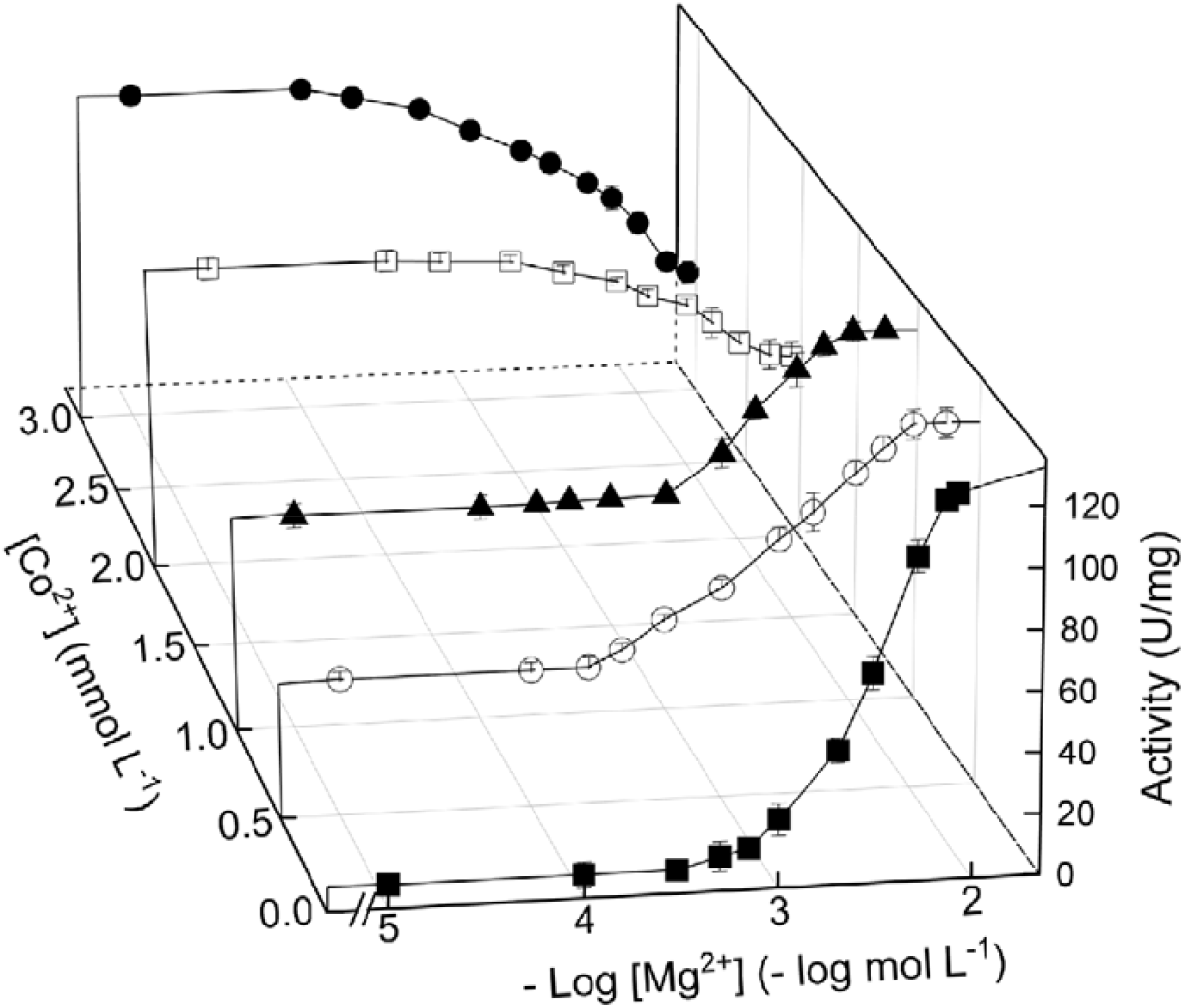
Effect of magnesium ions on the modulation of K^+^-phosphatase activity at different cobalt ion concentrations in *C. danae* gill (Na^+^, K^+^)-ATPase. Activity was assayed continuously at 25 °C in 50 mmol L^−1^ HEPES buffer, pH 7.5, containing 10 mmol L^−1^ *p*NPP, 15 mmol L^−1^ KCl and 9 µg alamethicin in a final volume of 1 mL. The mean activity of duplicate aliquots of the same microsomal preparation (≈15 µg protein) was used to fit the saturation curve which was repeated using three different microsomal preparations (± SD, N= 3). Where lacking, error bars are smaller than the symbols used. (◼) without Co^2+^. (◯) 0.5 mmol L^−1^ Co^2+^. (▴) 1 mmol L^−1^ Co^2+^. (□) 2 mmol L^−1^ Co^2+^. (•) 3 mmol L^−1^ Co^2+^.

### 3.4. Effect of Co^2+^ on K^+^ stimulation

Under saturating ionic and substrate concentrations (see legends to Fig. 5A and 5B) in the absence of Co^2+^, increasing K^+^ stimulated K^+^-phosphatase activity to a maximum rate of V_M_= 134.2 ± 4.5 nmol *p*NP^−^ min^−1^ mg^−1^ protein with K_0.5_= 9.60 ± 2.04 mmol L^−1^ (Fig. 5A and Table 1) following cooperative kinetics (n_H_= 2.2). Substitution of Mg^2+^ by 3 mmol L^−1^ Co^2+^ also gave a single saturation curve overlapping with that for Mg^2+^ and showing a maximum rate of V_M_= 133.3 ± 3.3 nmol *p*NP^−^ min^−1^ mg^−1^ protein with K_0.5_= 6.00 ± 1.50 mmol L^−1^ (Fig. 5A and Table 1). Ouabain-insensitive *p*NPPase activity was not stimulated by either metal ion over the concentration range used (inset to Fig. 5A). With Co^2+^ plus Mg^2+^, K^+^-phosphatase activity decreased to V_M_= 59.5 ± 4.0 nmol *p*NP^−^ min^−1^ mg^−1^ protein with K_0.5_= 2.79 ± 0.41 mmol L^−1^ following a single titration curve (Fig. 5B and Table 1), obeying cooperative kinetics (nH= 1.5). K^+^-phosphatase activity at K^+^ concentrations be low 10^−4^ mol L^−1^ was ≈15 nmol *p*NP^−^ min^−1^ mg^−1^ protein. Ouabain-insensitive *p*NPPase activity was stimulated to ≈22 nmol *p*NP^−^ min^−1^ mg^−1^ protein over the same K^+^ concentration range (inset to Fig. 5B). K^+^-phosphatase activity was not synergically stimulated by K^+^ plus NH_4_^+^ (not shown).

**Figure 5.**
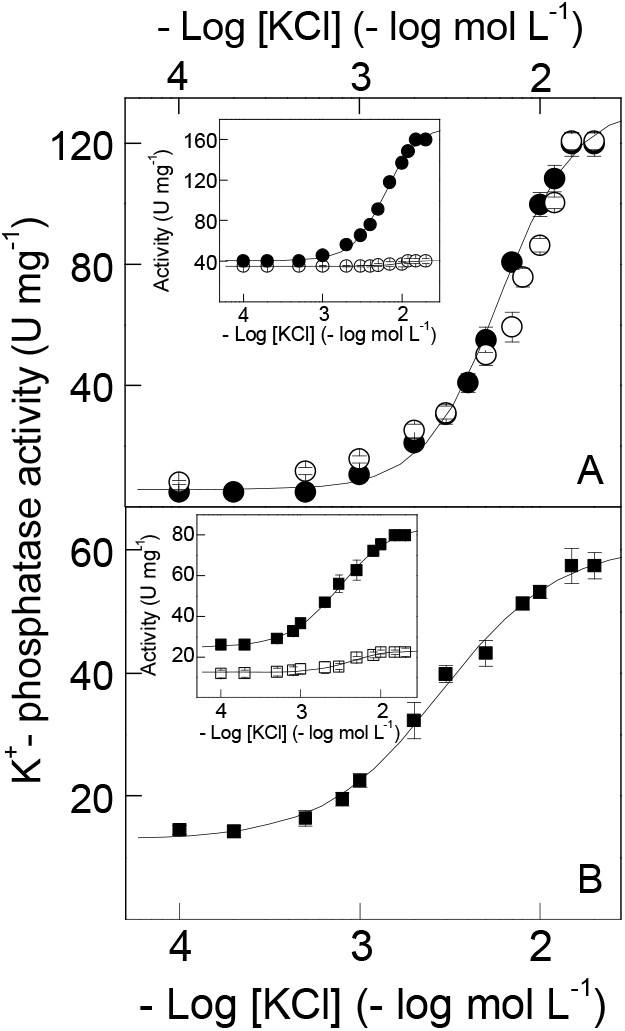
Effect of cobalt or magnesium ions on the modulation by potassium ions of K^+^-phosphatase activity of *C. danae* gill (Na^+^, K^+^)-ATPase. Activity was assayed continuously at 25 °C in 50 mmol L^−1^ HEPES buffer, pH 7.5, containing 10 mmol L^−^1 *p*NPP, 9 µg alamethicin and the metal ion (7 mmol L^−1^ Mg^2+^ or 3 mmol L^−1^ Co^2+^) in a final volume of 1 mL. The mean activity of duplicate aliquots of the same microsomal preparation (≈15 µg protein) was used to fit the saturation curve which was repeated using three different microsomal preparations (± SD, N= 3). Where lacking, error bars are smaller than the symbols used. **A-** K^+^-phosphatase activity with Mg^2+^ (◯) or Co^2+^ (•**). B-** K^+^-phosphatase activity with both Mg^2+^ and Co^2+^. Insets to figures**-** total *p*NPPase activity **( )** and ouabain-insensitive *p*NPPase activity (•) Co^2+^.

### 3.5. Effect of Co^2+^ on NH_4_^+^ stimulation

Under saturating ionic and substrate concentrations (see legends to Fig. 6A and 6B) in the absence of Co^2+^, a maximal rate of V_M_= 122.2 ± 5.2 nmol *p*NP^−^ min^−1^ mg^−1^ protein with K_0.5_= 9.02 ± 2.51 mmol L^−1^ was estimated for NH_4_^+^ concentrations increasing from 10^−4^ to 5×10^−2^ mol L^−1^ (Fig. 6A and Table 1). Substitution of Mg^2+^ by 3 mmol L^−1^ Co^2+^ also gave a single saturation curve with a maximum rate of V_M_= 127.9 ± 4.2 nmol *p*NP^−^ min^−1^ mg^−1^ protein with K_0.5_= 6.00 ± 1.10 mmol L^−1^, overlapping with that for Mg^2+^ (Fig. 6A and Table 1). Stimulation of ouabain-insensitive *p*NPPase activity by Mg^2+^ and Co^2+^ was negligible over the NH_4_^+^ concentration range used (inset to Fig. 6A). With Co^2+^ plus Mg^2+^, K^+^-phosphatase activity decreased to V_M_= 61.9 ± 3.7 nmol *p*NP^−^ min^−1^ mg^−1^ protein with K_0.5_= 5.46 ± 0.64 mmol L^−1^ (Fig. 6B and Table 1). Stimulation of the ouabain-insensitive *p*NPPase activity was <10% with Co^2+^ plus Mg^2+^ (inset to Fig. 6B). K^+^-phosphatase activity was not stimulated synergically by K^+^ plus NH_4_^+^ (not shown).

**Figure 6.**
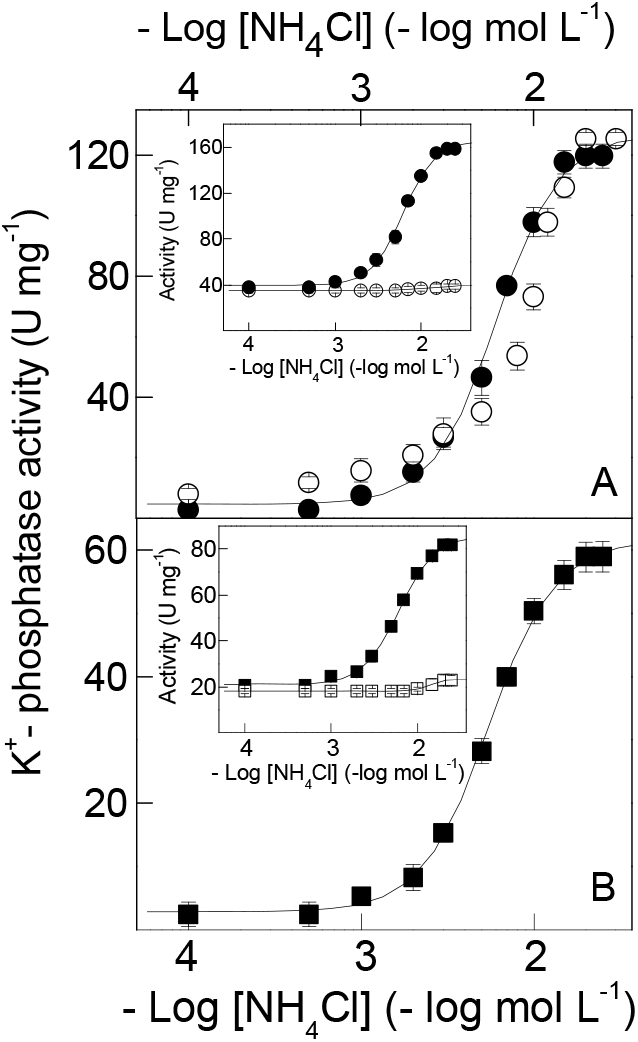
Effect of cobalt or magnesium ions on the modulation by ammonium ions of K^+^-phosphatase activity of *C. danae* gill (Na^+^, K^+^)-ATPase. Activity was assayed continuously at 25 °C in 50 mmol L^−1^ HEPES buffer, pH 7.5, containing 10 mmol L^−^1 *p*NPP, 9 µg alamethicin and the metal ion (7 mmol L^−1^ Mg^2+^ or 3 mmol L^−1^ Co^2+^) in a final volume of 1 mL. The mean activity of duplicate aliquots of the same microsomal preparation (≈15 µg protein) was used to fit the saturation curve which was repeated using three different microsomal preparations (± SD, N= 3). Where lacking, error bars are smaller than the symbols used. **A-** K^+^-phosphatase activity with Mg^2+^ (◯) or Co^2+^ (•**). B-** K^+^-phosphatase activity with both Mg^2+^ and Co^2+^. Insets to figures**-** total *p*NPPase activity (•) and ouabain-insensitive *p*NPPase activity (◯) Co^2+^.

### 3.6. Effect of Co^2+^ on inhibition by Na^+^ and ouabain

Cobalt does not affect inhibition of *p*NPPase activity by Na^+^ or ouabain (Fig. 7) Sodium concentrations from 10^−3^ to 0.2 mol L^−1^ inhibited *p*NPPase activity by ≈70% either with or without Co^2+^ (Fig. 7A and Table 1). The IC_50_ estimated for Na^+^ inhibition of *p*NPPase activity was ≈16 mmol L^−1^ (Table 1). Over the ouabain concentration range from 10^−6^ to 10^−2^ mol L^−1^, the Na^+^ and ouabain inhibition curves overlapped independently of Co^2+^ (Fig. 7B and Table 1). Their monophasic behavior with very similar inhibition constants (K_I_= 2.27 ± 0.62 mmol L^−1^ and K_I_= 2.13 ± 0.97 mmol L^−1^ for Mg^2+^ and Co^2+^ respectively) (inset to Fig. 7B) suggests a single ouabain binding site.

**Figure 7.**
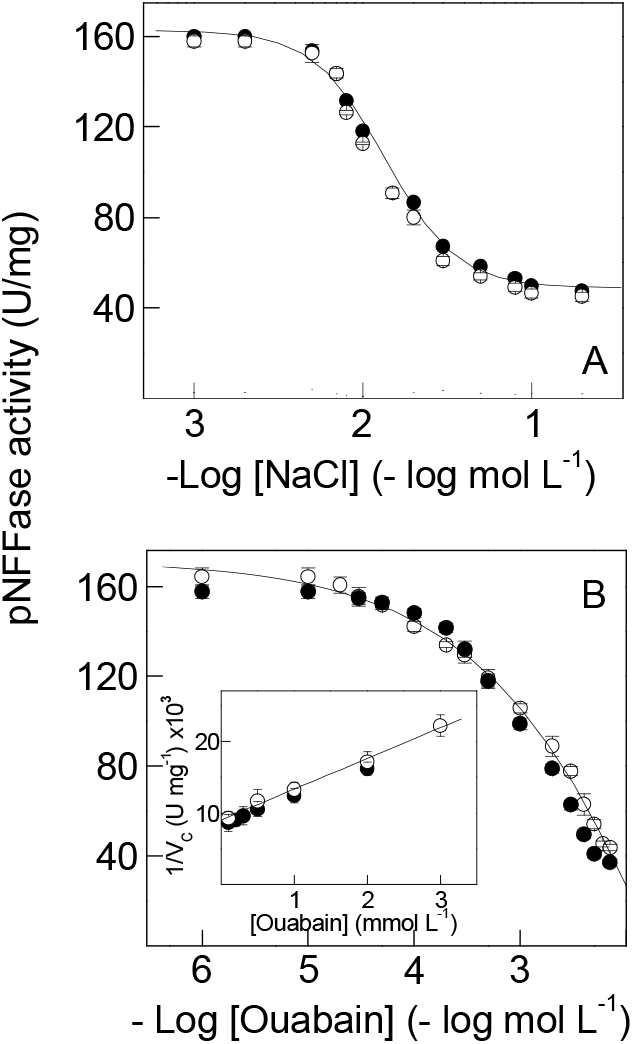
Effect of cobalt or magnesium ions on the inhibition by sodium and ouabain on *p*NPPase activity of *C. danae* gill (Na^+^, K^+^)-ATPase. Activity was assayed continuously at 25 °C in 50 mmol L^−1^ HEPES buffer, pH 7.5, containing 10 mmol L^−1^ *p*NPP, 15 mmol L^−1^ KCl and 9 µg alamethicin in a final volume of 1 mL. The mean activity of duplicate aliquots of the same microsomal preparation (≈15 µg protein) was used to fit the saturation curve which was repeated using three different microsomal preparations (± SD, N= 3). Where lacking, error bars are smaller than the symbols used. **A-** sodium ion. **B-** ouabain. Inset to Fig. 8B-Dixon plot for estimation of K_I_ in which v_c_ is the *p*NPPase activity corrected for residual *p*NPPase activity found at high ouabain concentrations. (○) with 7 mmol L^−1^ Mg^2+^. (•) with 3 mmol L^−1^ Co^2+^.

## 4. DISCUSSION

We provide a comprehensive analysis of the effects of Co^2+^ on the modulation *in vitro* of the K^+^-phosphatase activity in a gill (Na^+^, K^+^)-ATPase from the blue crab *Callinectes danae*. Depending on Mg^2+^, Co^2+^serves as both stimulator and inhibitor of K^+^-phosphatase activity. Without Mg^2+^, Co^2+^ stimulates activity as does Mg^2+^, although with a ≈4.5-fold greater affinity. With Mg^2+^, activity is almost completely inhibited by Co^2+^, while ouabain inhibition is unaffected. Substitution of Mg^2+^ by Co^2+^ slightly increases enzyme affinity for K^+^ and NH_4_^+^. Mg^2+^ displaces bound Co^2+^ from the Mg^2+^-binding site in a concentration dependent manner; however, Co^2+^ does not displace bound Mg^2+^ even at elevated concentrations. Saturation by Co^2+^ of the Mg^2+^-binding site does not affect substrate recognition by the enzyme.

K-phosphatase activities estimated with Mg^2+^ or Co^2+^ are similar and their overlapping *p*NPP saturation curves show comparable cooperative effects; their similar K_0.5_ values suggest that Co^2+^ saturation of the Mg^2+^-binding site does not affect substrate recognition. The high stability constants for the Co^2+^-*p*NPP (130 mol L^−1^, [76]) and Mg^2+^-*p*NPP (170 mol L^−1^, [77]) complexes suggest that negligible free metal ions are present at millimolar metal ion concentrations, i.e., the metal-*p*NPP complex is the true enzyme substrate. The lower apparent dissociation constant for Co^2+^ (K_0.5_= 1.15 ± 0.61 mmol L^−1^), close to that for Mg^2+^ (K_0.5_= 1.76 ± 0.49 mmol L^−1^), is comparable to the enzyme from *Cancer pagurus* axonal membranes despite its 2-fold greater maximum *p*NPP hydrolysis rate [78].

Millimolar Mg^2+^ concentrations are required for K^+^-phosphatase activity of the *C. danae* (Na^+^, K^+^)-ATPase and, like the mammalian enzyme [58,79], no detectable activity can be measured without this ion. The millimolar Mg^2+^ or Co^2+^ concentrations necessary for maximum K^+^-phosphatase activity exclude the likelihood of metal binding other than Mg^2+^ or Co^2+^ to the Mg^2+^-binding site during the catalytic cycle [56]. The inhibition by free Mg^2+^ or Co^2+^ of K^+^-phosphatase activity may result from competition with K^+^ for the Mg^2+^-binding site [56,61,80] or to excess Mg^2+^ bound during the E2K conformation, decreasing affinity for *p*NPP [52,56,81].

Co^2+^ can substitute for Mg^2+^, stimulating K^+^-phosphatase activity more efficiently (K_0.5_ ≈4.5-fold lower). However, Co^2+^ does not displace Mg^2+^ from the Mg^2+^-binding site of the *C. danae* enzyme. Inhibition by excess Co^2+^ likely results from Co^2+^ binding at a site different from the Mg^2+^-binding site. Two distinct Mg^2+^ binding sites are known for crustacean [73] and mammalian [50] (Na^+^, K^+^)-ATPases, although only one is relevant for *p*NPPase and ATPase activities [82]. Like Co^2+^, Mn^2+^ also stimulates sheep kidney (Na^+^, K^+^)-ATPase activity, the K_D_= 0.88 μmol L^−1^ for Mn^2+^ binding being very similar to the kinetic constant for ATP hydrolysis [83]. Co^2+^ stimulates dog kidney outer medulla (Na^+^, K^+^)-ATPase activity by substituting for Mg^2+^ [20]. Differently from Co^2+^, Cu^2+^ inhibits the gill (Na^+^, K^+^)-ATPase activity of rainbow trout *Oncorhynchus mykiss* [84], and the K^+^-phosphatase and (Na^+^, K^+^)-ATPase activities of rabbit kidney [85] by directly interfering with Mg^2+^ binding, affecting Mg•ATP hydrolysis.

The similar V_M_ and K_0.5_ values for stimulation by K^+^ or NH_4_^+^ of K^+^-phosphatase activity with Mg^2+^ or Co^2+^ suggests that NH_4_ ^+^ binds to the same site as K^+^ during the catalytic cycle, independently of Mg^2+^ or Co^2+^ bound to the Mg^2+^-binding site. While K^+^-phosphatase activity is inhibited to same degree by Mg^2+^ and Co^2+^, the 2-fold greater K_0.5_ for NH_4_^+^ (5.46 mmol L^−1^) compared to K^+^ (2.79 mmol L^−1^) suggests that Co^2+^ binding to a different Mg^2+^-binding site affecting enzyme interaction with NH ^+^. Without Na^+^, stimulation by K^+^ of K^+^-phosphatase activity involves two K^+^ binding sites: one that regulates *p*NPP access to the phosphatase site, the other increasing catalytic activity [86]. ATP binding to the low-affinity substrate binding site induces a conformational change in the cytoplasmic domain of the enzyme attributed to the E2 to E1 transition; the subsequent binding of Mg^2+^ to the enzyme ATP complex induces a new conformational change that facilitates the E1 to E2 transition [87]. Like Mg^2+^, Co^2+^ may induce a conformational change, stimulating the enzyme during *p*NPP hydrolysis.

K^+^-phosphatase activity can be stimulated by Tl^+^, Rb^+^ or NH_4_ ^+^ to rates similar to K^+^ stimulation while Cs^+^ and Li^+^ exhibit lower stimulation (5-30%) [88,89]. For 21‰ (low salinity)-acclimated *C. danae*, Rb^+^ stimulates gill *p*NPPase activity by 1.5-fold compared to K^+^ [63]. NH_4_^+^ may sustain ATP hydrolysis by replacing K^+^ [89,90] and is actively transported by crustacean and vertebrate enzymes [91,92]. Together with K^+^, NH_4_^+^ synergically stimulates ATP hydrolysis by the *C. danae* (Na^+^, K^+^)-ATPase through an additional increment, strongly influenced by Mg^2+^ and Na^+^, underlying ammonia excretion in crustaceans [90]. The E2 conformation is the main state responsible for *p*NPP hydrolysis [58] and is likely the reason that K^+^-phosphatase activity is not synergically stimulated by K^+^ and NH_4_^+^. Synergic stimulation using *p*NPP as a substrate is known exclusively for the shelling crab *C. ornatus* [63]. When using ATP as a substrate, species-specific synergic stimulation is found in various crustaceans [73].

The inhibition by Na^+^ of *C. danae p*NPPase activity independently of Mg^2+^ or Co^2+^ is seen also for (Na^+^, K^+^)-ATPases from various sources [73], and reflects competition by Na^+^ for the cytoplasmic K^+^-binding sites, favoring the E1 conformation [50,56,58,93–95]. Na^+^ inhibition also may involve events other than the simple binding of the ion. To illustrate, the synergistic stimulation by Na^+^ and K^+^ (3%) of *p*NPPase activity in the electric organ of *Electrophorus electricus* [86] suggests that both ions bind to different sites. Both 10 mmol L^−1^ Na^+^ and 15 mol L^−1^ K^+^ inhibit the *C. danae p*NPPase activity by ≈40%, as seen in the freshwater shrimp *Macrobrachium olfersii* [61] and crabs *Cancer pagurus* [95] and *C. ornatus* [63] under identical assay conditions. Na^+^ inhibits *p*NPPase activity allosterically in various mammalian and crustacean (Na^+^, K^+^)-ATPases [61,63,86,89,95] including *M. olfersii* [61] and *C*. *pagurus* [95]. Thus, considering a two K^+^-binding site model [86], at low (10-fold less than K^+^) concentrations, Na^+^ competes for the high-affinity K^+^ binding site, leading to allosteric effects. At high concentrations (similar to K^+^), Na^+^ competes for the low-affinity K^+^-binding site, reducing maximum hydrolysis rate.

Co^2+^ does not affect ouabain binding to the *C. danae* gill (Na^+^, K^+^)-ATPase, the single inhibition curve and K_I_ being very similar to Mg^2+^. Most species, except the red river crab *Dilocarcinus pagei* [96], exhibit a single ouabain inhibition curve independently of substrate [62,63,72,97,98]. The K_I_ for ouabain inhibition with Co^2+^ lies in the range for various (Na^+^, K^+^)-ATPases [73]. It should be noted that only for the cerebromicrovascular (Na^+^, K^+^)-ATPase, Pb^2+^ and Al^3+^ caused selective alterations in ATP hydrolysis inhibiting and stimulating ouabain binding, respectively [99].

Our findings reveal that without Mg^2+^, Co^2+^ stimulates the gill (Na^+^, K^+^)-ATPase to levels similar to Mg^2+^. Without Mg^2+^, Co^2+^ stimulates K^+^-phosphatase activity similarly although with a ≈4.5-fold greater affinity than with Mg^2+^, which almost completely inhibits K^+^-phosphatase activity. Mg^2+^ displaces Co^2+^ from the Mg^2+^-binding sites in a concentration dependent manner. Ouabain inhibition is identical with Co^2+^ or Mg^2+^. Saturation by Co^2+^ of the Mg^2+^-binding sites does not affect substrate recognition by the enzyme. Given the complex interactions between heavy metal ion contaminants and enzymes, their toxic effects at the molecular level are poorly understood. Our findings contribute to elucidate partly the mechanism of action of Co^2+^ on a crustacean gill (Na^+^, K^+^)-ATPase.

## Acknowledgements

The authors thank the Instituto Chico Mendes de Conservação da Biodiversidade, Ministério do Meio Ambiente (ICMBio/MMA) for authorizing collecting permit #29594-18 to JCM, and INCT-ADAPTA II (Instituto Nacional de Ciência e Tecnologia para Adaptações da Biota Aquática da Amazônia, ADAPTA-II) with which FAL’s laboratory is integrated, and the Amazon Shrimp Network (Rede de Camarão da Amazônia).

## Funding information

This investigation was financed by research grants from the Fundação de Amparo à Pesquisa do Estado de São Paulo (FAPESP 2016/25336-0 and 2019/21899-8), Fundação de Amparo à Pesquisa do Estado de Minas Gerais (FAPEMIG APQ-01893-16), Conselho de Desenvolvimento Científico e Tecnológico (CNPq 458246/2014-0), and in part by INCT ADAPTA II (CNPq 465540/2014-7) and the Fundação de Amparo à Pesquisa do Estado do Amazonas (FAPEAM 062.1187/2017).

FAL (302072/2019-7), and JCM (303613/2017-3) received Excellence in Research scholarships from CNPq. LMF, CMM and MICC received scholarships from the Coordenação de Aperfeiçoamento de Pessoal de Nível Superior (CAPES, Finance code 001).

## Author contributions

**Francisco A. Leone**: Conceptualization, Formal analysis, Resources, Funding acquisition, Methodology, Supervision, Project administration. Writing original draft, Review & Editing. **Leonardo M. Fabri**: Methodology, Investigation, Conceptualization, Writing original draft, Review & Editing. **Cintya M. Moraes**: Methodology, Investigation, Writing original draft. **Maria I. C. Costa**: Methodology, Investigation, Writing original draft. **Daniela P. Garçon**: Conceptualization, Methodology, Formal analysis, Funding acquisition, Writing original draft, Review & Editing. **John C. McNamara**: Conceptualization, Methodology, Formal analysis, Writing original draft, Review & Editing.

## Data availability

The datasets generated and/or analyzed during this study are available from the corresponding author on reasonable request.

## Declaration of competing interests

All authors certify that they have no affiliations with or involvement in any organization or entity with any financial or non-financial interest in the subject matter or materials discussed in this manuscript.

## Ethical approval studies in animals

This investigation complies with all local, state, federal and international guidelines as regards the use of invertebrate animals in scientific research. This study also complies with the ARRIVE guidelines.

## Notes

### Competing Interest Statement

The authors have declared no competing interest.

